# Asgard archaea defense systems and their roles in the origin of eukaryotic immunity

**DOI:** 10.1101/2023.09.13.557551

**Authors:** Pedro Leão, Mary E. Little, Kathryn E. Appler, Daphne Sahaya, Emily Aguilar-Pine, Kathryn Currie, Ilya J. Finkelstein, Valerie De Anda, Brett J. Baker

## Abstract

Immune systems are integral to survival against viral infection. Recently, dozens of new antiviral systems have been characterized in bacteria^1^. Some of these systems are present in eukaryotes and appear to have originated in prokaryotes. However, little is known about these defense mechanisms in archaea. Here, we identified 2,610 complete defense systems in archaea related to eukaryotes, the Asgardarchaeota^2^. These comprise 89 unique systems, including argonaute, NLR, mokosh, viperin, lassamu, and CBASS. Asgard viperin (asVip) and argonaute (asAgo) proteins are present at high frequencies compared to bacteria and have structural homology to eukaryotes. Phylogenetic analyses revealed asVips are ancestral eukaryotic proteins. Heterologous expression of asVips in bacteria, including the lineage closest to eukaryotes, Hodarchaeales, showed anti-phage activity. Eukaryotic- and bacterial-argonaute proteins appear to have originated in Asgardarchaeota and preserve ancient structural characteristics. asAgos have argonaute-PIWI domains which are key components of the RNA interference (RNAi) in eukaryotes. Characterization of hundreds of defense systems in the Asgardarchaeota revealed these archaea played important roles in the innovation of viral protection in eukaryotes. Given their relationship to eukaryotes, these defense systems may have applications in biomedicine and biotechnology.

---

Organisms across the tree of life contain complex defense systems (DS) to battle viral infections^3,4^. Over the past decade, dozens of new DS have been identified and characterized in bacteria, sparking a debate about a potential link between these systems and the origins of innate immune mechanisms in eukaryotes. More recently, protein components of bacterial NLR (Nucleotide-binding domain leucine-rich repeat), CBASS (Cyclic oligonucleotide-based antiphage signaling system), viperins (virus-inhibitory protein, endoplasmic reticulum-associated, interferon (IFN)-inducible), argonautes, and other DS have been shown to exhibit homology with proteins involved in the eukaryotic immune system^5^. Most of the research on prokaryotic defense systems has focused on bacteria, with archaea representing <3% of the genomes in these studies^6,7,8^. Thus, very little is known about the diversity or evolution of these systems in archaea.

Recently, diverse novel genomes have been obtained belonging to the archaea most closely related to eukaryotes, commonly referred to as “Asgard” archaea, the phylum Asgardarchaeota^2^. In addition to being sister lineages to eukaryotes, these archaea also contain an array of genes that are hallmarks of complex cellular life involved in signal processing, transcription, translocations, among other processes^9^. The Asgard archaea are descendants of the ancestral host that gave rise to eukaryotic life. One newly described order, the Hodarchaeales (within the Heimdallarchaeia class), shared a common ancestor with eukaryotes^2^. Here, we characterize defense systems in archaea and show that Asgard archaea have a broad array of these DS. We also show that Asgards contributed to the origins of innate immune mechanisms in eukaryotes.

To explore the diversity and distribution of DS in archaea, we used 132 previously described defense systems to search through a comprehensive dataset containing 3,408 publicly available genomes, and a newly expanded set of Asgardarchaeota genomes (869). A total of 30,761 defense system-associated genes have been identified across these archaeal genomes, belonging to 11,466 complete defense systems, with an average of 4.55 DS genes per archaeal genome (Figure 1; Supplementary Data 1). Of a total of 132 previously described DS, 89 were found in these genomes. 27.5% of these archaeal genomes do not contain any known DS. A total of 2,610 complete DS were identified in Asgardarchaeota genomes. These varied among the classes between 1 and 65, with an average of 4.9 DS per genome. These numbers are similar to those reported in bacteria, on average 5.6 DS identified by Tesson *et al*.^7^ and 5.8 DS by Millman *et al*.^8^. The DPANN archaea contain the lowest number, with a mean of 2.6 DS per genome. As previously reported in bacteria, the most prevalent DS in archaea are the restriction-modification (RM) and CRISPR systems^7,10^. These DSs represent 45% and 22% of all known defense systems in archaeal genomes, respectively (Supplementary Data 1).

**Figure 1.**
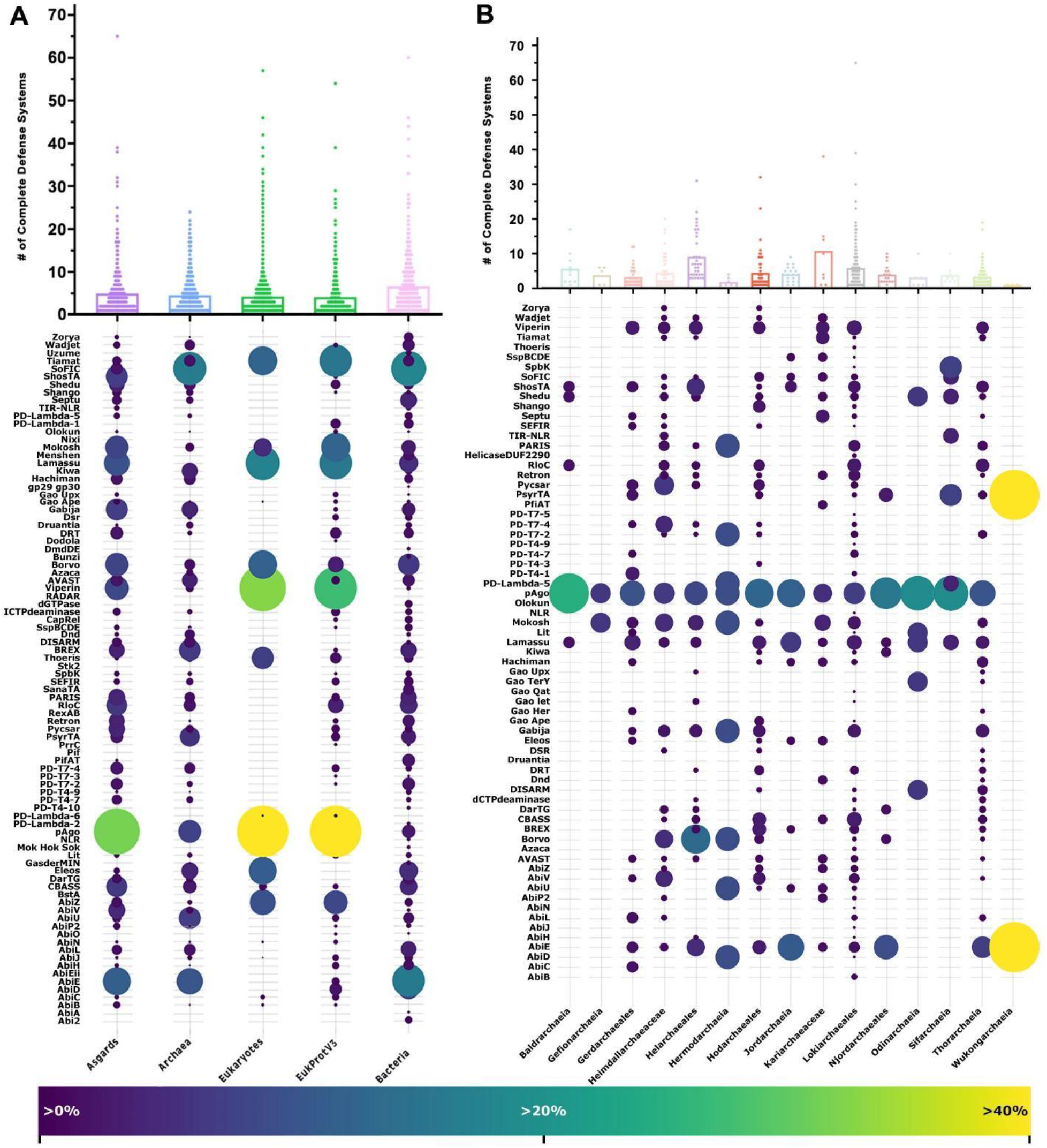
Distribution of defense systems across the tree of life and in Asgard archaea. **A**) The total number of complete defense systems (DS) in each genomic dataset used in this study. The bubble plot shows the frequency of distribution of DS identified (Y-axis) in the datasets (X-axis). **B)** The total number of DS across Asgardarchaeota genomes. Bubble plot: frequency of distribution of DS in each Asgardarchaeota group. Dots in the graphs represent the number of DS in a single genome from the respective group. The size and the color of the dots in the bubble plot are proportional to the prevalence of the DS in that group. For clarity, the 2 most prevalent DS (CRISPR and RM) across the genomic datasets were removed from both bubble plots representation. DS less found in Eukaryotes (check Methods section for the list) were also removed from the bubble plot in panel A.

Asgardarchaeota have a broad array of DS compared to those in bacteria. Some Asgard archaeal groups have a higher DS per genome ratio compared to other archaea (Baldrarchaeia 5.7), and others surpass bacteria (Helarchaeales 9.1; Kariarchaeaceae 10.7; Lokiarchaeles 5.8). Four out of the five representatives from the Heimdallarchaeia class, including genomes from Hodarchaeales, have fewer DS (Njordarchaeales 4.0; Gerdarchaeales 3.3; Heimdallarchaeaceae 4.5; Hodarchaeales 4.4) (Figure 1B; Graph), suggesting they have yet uncharacterized DS. Given that Heimdallarchaeia is the prokaryotic class most closely related to eukaryotes^2^, it is possible that these organisms might utilize mechanisms more similar to those observed in unicellular eukaryotes that were not detected using a prokaryotic DS database.

Twenty-two DS are more frequently found in Asgardarchaeota genomes compared to other prokaryotes (AbiB, AbiP2, AbiV, Argonautes, Borvo, Cas, dCTPdeaminase, Gao Ape, Gao Her, Gao Iet, hachiman, pycsar, PD-T7-4, PD-T4-1, PD-T4-7, RloC, Rst helicase, Rst2TM TIR-NLR, ShosTA, Shango, Shedu, viperins). Conversely, 12 DS are more abundant in archaea from groups outside of Asgardarchaeota (Figure 2B; Bubble plot). Viperins and argonautes account for 2.1% and 7.9% (respectively) of all DS identified in Asgardarchaeota genomes. Viperins and argonautes are found 9 and 4 times more frequently in Asgard genomes than in other archaeal groups. When compared to bacteria, the disparity is even more pronounced, with 29 and 11 times higher representation of viperins and argonautes in Asgardarchaeota. Argonautes are present in every Asgardarchaeota class.

**Figure 2.**
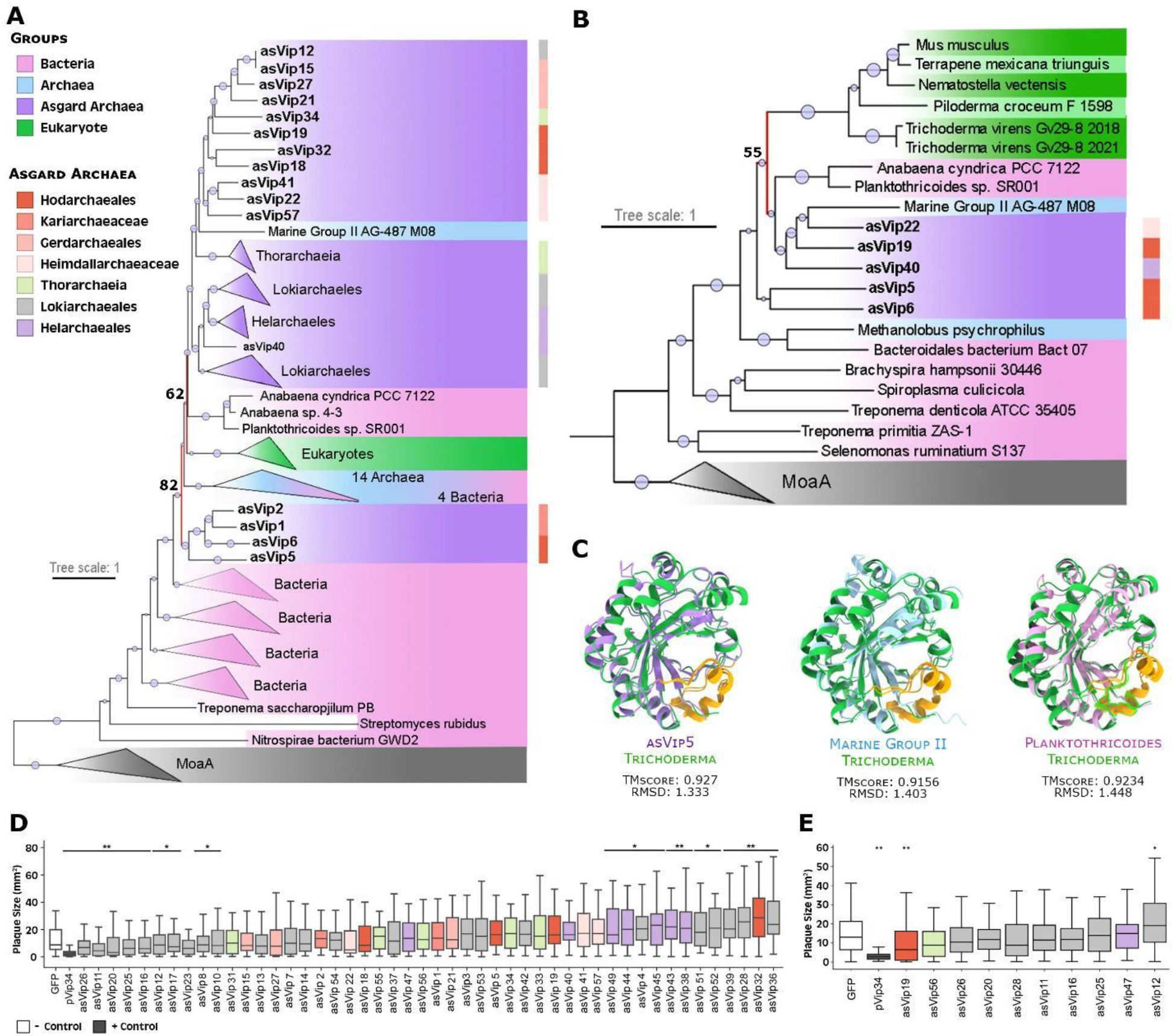
Evolutionary history and anti-phage activity of Asgard viperins. **A)** Phylogenetic analysis of viperins. Viperins phylogeny revealed ancestral links of eVip (eukaryotic viperin) with asVip (asgard viperin) (nodes marked in red), particularly those within the Heimdallarchaeia class (including Kariarchaeaceae (2), Heimdallarchaeaceae (3) and Hodarchaeales (5)). The size of the dots on the nodes is proportional to bootstrap values ranging between 60 and 100. **B)** Structure-based homology of viperins. Consistent with the sequence homology-based phylogenetic tree, the eVip structure appears to have been inherited from asVip (red node). The darker green color represents reference sequences predicted experimentally. The size of the dots at the center of the nodes is proportional to bootstrap values ranging between 50 and 100. **C)** Superposition of an eVip structure, predicted by X-ray diffraction (green), and the structural models of an asVip, archaeal viperin (arVip), and bacterial viperin (from left to right). The yellow color in the models emphasizes the high conservation of the viperin catalytic site across the tree of life. **D)** Anti-T7 phage activity of asVip in *E. coli*. Nine asVip (asVip 26,11,20,25,16,12,17,23,8) exhibited anti-viral activity as indicated by the p-values (* p < 0.05; ** p < 0.01). **E)** Anti-T7 phage activity of asVip after codon optimization for their expression in *E. coli*. One asVip from a Hodarchaeales organism provided protection against viral infection (asVip 19). The center line of each box plot denotes the median; the box contains the 25^th^ to 75^th^ percentiles. Black whiskers mark the 5^th^ and 95^th^ percentiles. pVip34 is a prokaryotic viperin selected as a positive control from Bernheim *et al*., 2021^13^. Each experimental condition includes, on average, 53 plaques pooled from three biological replicates.

Viperins were first described in eukaryotes as one of the key players in the mechanism of inhibition of human cytomegalovirus (HCMV) infection^11^ and later found in prokaryotes^4^. Viperin proteins in archaea and bacteria have sequence and structural residues conservation^12^. These structural residues are a strong indicator of a conserved defense mechanism^13^. Protein sequence and structural conservations make it possible to reconstruct protein phylogenies, which indicate a prokaryotic origin and show Asgard archaea as being a sister group to eukaryotes, suggesting eukaryotic and Asgard viperins evolved from a common protein ancestor (Figure 2).

Using an expanded archaeal genomic dataset our phylogenies revealed eukaryotic viperins (eVip) and Asgard viperins (asVip) are sister proteins and share a common ancestor (Figure 2A). Only four proteins outside the Asgard group (one archaeal and three bacterial) are present in this clade. The two bacterial species harboring these viperin sequences are cyanobacteria belonging to the genera *Anabaena* and *Planktothricoides*, known for their symbiotic relationships with eukaryotic organisms. *Anabaena* spp. are present throughout the entire lifecycle of plants from the *Azolla* genus^14^. Co-speciation and gene transfer have been reported in these host-symbiont interactions^15^, which could explain the position of these viperins near eukaryotes. The diversity of basal asVip to eukaryotic sequences suggests that this immune mechanism in eukaryotes was derived from Asgardarchaeota (Figure 2A, red nodes).

Viperin proteins are structurally conserved across all domains of life (Figure 2B). As a member of the radical S-adenosylmethionine superfamily, the SAM domain is present in all viperins characterized by a partial (βα)6-barrel folds located at the center of the protein^16^. The most conserved structure is the catalytic site cavity closer to the C-terminal extension. This site is present in asVips (yellow in Figure 2C), indicating that the 3′-deoxy-3′,4′-didehydro (ddh) synthase activity has been preserved among all domains of life (Figure 2C). Concomitant with our study, Shomar et al. provided experimental support for this proposed conservation^17^. These findings provide evidence for its conserved function as a defense mechanism, since ddh is the primary component of this DS, acting as the chain terminator in viral DNA/RNA polymerase in eukaryotes^18^, and prokaryotes^13^, and potentially in Asgardarchaeota (Figure 2D and E; discuss below).

To ensure that asVips can protect cells against viral infections, we challenged asVip-expressing bacteria with T7 phage. Both bacterial and eVips have previously been demonstrated to inhibit this phage’s infection^13^. We synthesized and constitutively expressed 48 Asgard viperins in *E. coli*, which were then submitted to testing (Supplementary Data 2). At first, 9 out of the 48 asVips (19%), all belonging to Lokiarchaeales group, displayed anti-T7 defense activity (Figure 2D).

In a second assay, *asVip* were redesigned with codon optimization for heterologous expression in *E. coli*. Remarkably, a viperin originating from a Hodarchaeales genome was then capable of safeguarding cells against T7 phage infection (Figure 2E). Asgard archaea are known for possessing ribosomal architecture akin to eukaryotes^19^. More recently, representatives from Hodarchaeales have been found to contain a homologue of the previously eukaryote-exclusive ribosomal protein L28e^2^. The lack of viral protection observed in our initial assay could be indicative of the inefficiency of bacterial systems in expressing Asgard viperins from groups more closely related to eukaryotes, such as Hodarchaeales. This obstacle was overcome following codon optimization, which allowed the protein to be expressed more efficiently and demonstrate its effectiveness in protecting cells against viral infection. These analyses show that asVips are consistent with the defensive roles against viral infection observed for viperins previously described in both bacterial and eukaryotic contexts^13,18^.

In eukaryotes, both microRNAs (miRNAs) and small interfering RNAs (siRNAs) serve as guides that target RNA transcripts for degradation by the RNA-induced silencing complex (RISC) as a form of post-transcriptional regulation and cellular defense^20^. The signature proteins of this complex are the argonautes^21^. These proteins are members of the PIWI (P element–induced wimpy testis) superfamily, distinguished by the presence of the homonyms domain^22^. Argonaute-PIWI proteins are key components of the RNA interference (RNAi) system in eukaryotes. Prokaryotic argonautes (pAgos) are also associated with defense against mobile genetic elements^7,23,24,25^. pAgos are classified into long pAgos and short pAgos. Long pAgos domains organization is similar to that of eukaryotic argonautes (eAgos), comprising a N-terminal domain that serves as a wedge to separate the guide oligonucleotide from its target, a MID (middle) domain, and a PAZ (PIWI-Argonaute-Zwille) domain that together, anchors and stabilizes the guide molecule. Long pAgos also have a catalytic PIWI domain responsible for the cleavage of the target DNA/RNA^26,27^. Short pAgos possess only the MID and an inactive PIWI domain^24^.

Previously, the origin of eAgo has been proposed to be from euryarchaeal argonautes, and an ancient phylogenetic split between short and long types of these proteins was observed^23,24,28^. Phylogenetic analysis of archaeal argonautes (arAgo) identified in this study initially supported the early bifurcation into short and long forms (Supplementary Figure 1). However, it also revealed that Asgardarchaeota argonautes (asAgo) are the likely precursor of eAgo (Figure 3A, red nodes; Supplementary Figure 1). Our phylogenetic analysis also reveals a deep-rooted clade of asAgo as the ancestral form of long bacterial and eukaryotes argonautes (Figure 3A). An early clade of asAgo diversified into a larger group that encompasses arAgo and more than 98% of eAgo (Figure 3A). Interestingly, we also found that each clade of long proteins across the phylogeny is rooted by asAgos to some extent. This topological arrangement lends strong support to the hypothesis that an early diversification of asAgo gave rise to the wide array of argonaute proteins observed across the tree of life, with occasional instances of horizontal gene transfer (HGT).

**Figure 3.**
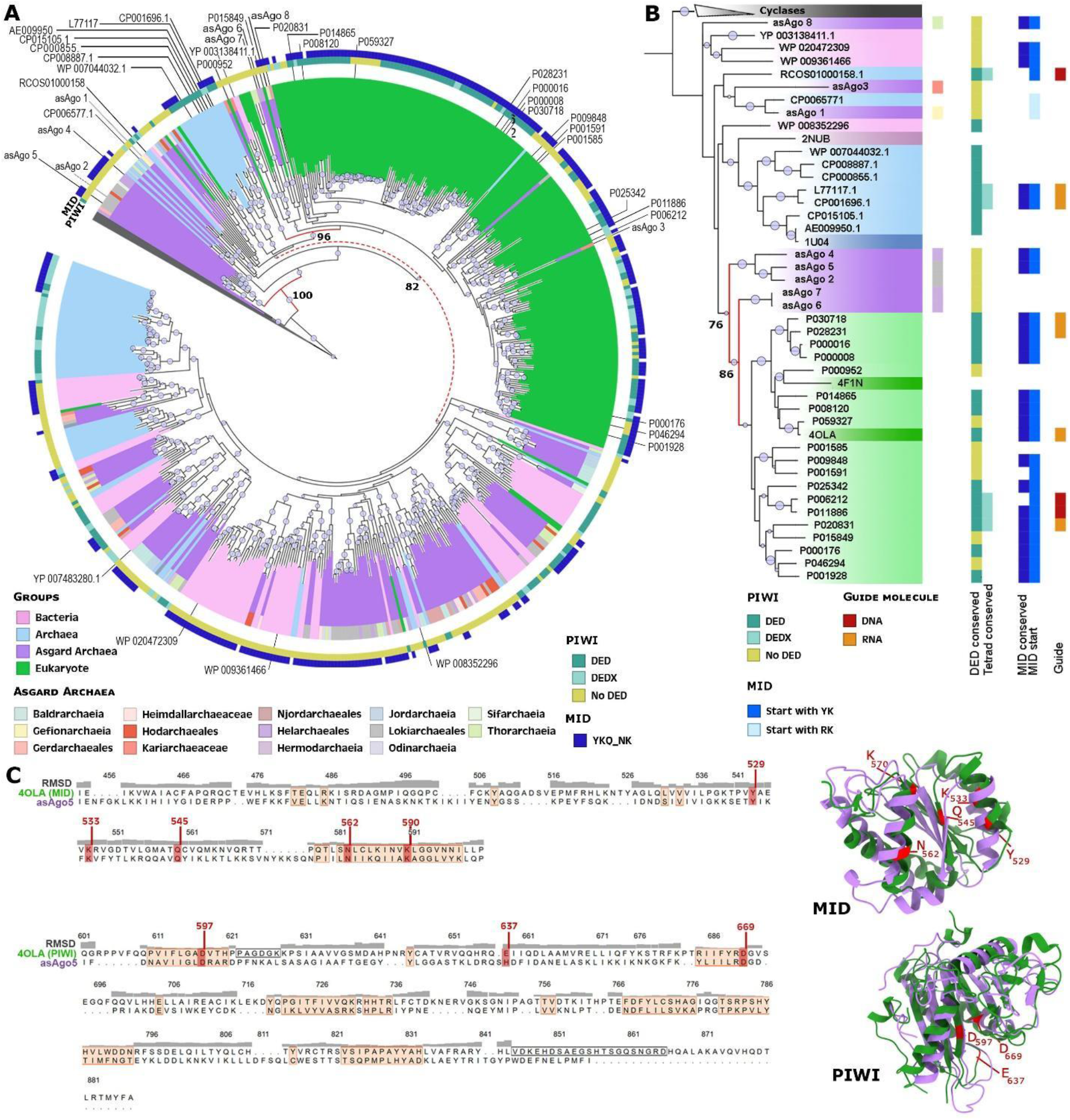
Evolutionary history of Asgard argonaute proteins. **A)** Phylogeny of long type argonaute proteins from archaea, bacteria, and eukaryotes with cyclases as outgroup (grey). **B)** Structure-based homology of argonautes. **C)** Structural alignment of asAgo5 and 4OLA (eAgo) MID and PIWI domains (left), and the graphic model of the corresponding alignments (right). Salmon regions on the alignment highlight strong conservation (low RMDS values). Red amino acids in the structural alignment, and their respective models represent the 4OLA conserved functional residues in MID and PIWI.

The predicted structure of argonautes revealed a clear conservation between asAgo and eAgo (Figure 3B; red nodes). Detailed examination of five asAgo most closely related to eAgo revealed that three of the proteins initially identified as long types are actually short Agos due to the absence of the PAZ domain (asAgo 2, 4, and 5). The remaining two (asAgo 6 and 7) possess only an isolated PIWI domain, a feature previously reported in approximately 1% of short Agos^28^. Intriguingly, four of these five proteins (asAgo 2, 4, 6, 7) contain a region at the N-terminal portion that is homologous to the eukaryotic translation initiation factor 2 (eIF2). This protein is a member of the PIWI superfamily and is seldom found in prokaryotes. eIF2 plays a critical role in delivering methionyl-tRNA to the 40S ribosomal subunit and subsequently binding to the 5’ end of capped eukaryotic mRNAs in conjunction with other eIFs^29,30^. Bioinformatics analysis shows a higher occurrence of eIF2 sequences in asAgo compared to other pAgos (Supplementary Data 4). This suggests the possibility of a ribosomal architecture and eukaryotic-like translation processing in these archaea. This is a hypothesis that still needs experimental confirmation.

Only 20% of the asAgo proteins contain DED catalytic triads, previously thought to be essential for PIWI domains to execute their nuclease function^31^. Of these asAgos (32 in total), 28% have the full tetrad DEDX of amino acids (X = D, H, N or K) (Figure 3; green and yellow squares) recently shown to be the set of residues necessary for maintaining PIWI activity^32^. These numbers are in accordance with the overall percentage of long pAgo predicted to be active. However, these findings challenge the initial assumption that the ancestral pAgo was an active nuclease^33^.

Structural alignment between asAgo and eAgo reveals a high degree of conservation in the PIWI and MID protein domains, even among those asAgos lacking a conserved PIWI tetrad (Figure 3C). This suggests that ancient asAgo enzymes contained all the components necessary for PIWI nuclease function but lacked the catalytic site. This pre-existing conserved protein organization likely facilitated the perpetuation of this basic structure over time, paving the way for its evolution toward an active nuclease enzyme. Support for this hypothesis comes from the higher number of asAgos with a conserved MID domain binding pocket (YKQ-NK). Experimental evidence has shown that the presence of tyrosine as the first amino acid in the pocket is crucial for a stable binding of the guide nucleotide to argonautes. This is due to threonine’s aromatic ring, which functions as a cap to the first base on the 5’ end of the guide molecule^28^.

Recently, experimental evidence revealed the function and mechanism of action of an asAgo from a Lokiarchaeales genome. Bastiaanssen *et al*. characterized the RNA-guided/RNA silencing capabilities of asAgo, suggesting it as a possible origin of this eukaryotic feature before eukaryogenesis^34^. This serves as a notable example of early functionalization of asAgo, where a new function emerges through the adaptation of pre-existing conserved domains.

The single-protein DS known as NLR, named after mammalian Nod-like receptors, serves as an intracellular sensor through its NACHT module, which is responsible for recognizing specific molecular signatures. Recent research suggests that eukaryotes acquired NLR via horizontal gene transfer from bacterial sequences^35^. Our phylogenetic analysis revealed a correlation between the NLR proteins in archaea and eukaryotes (Supplementary Figure 2A), suggesting they share a common ancestor. Moreover, certain groups of eukaryotic NACHT proteins appear to have a shared ancestry with Asgardarchaeota proteins (Supplementary Figure 2A; red node). However, more comprehensive analyses and additional NLR-related proteins are required to enhance the resolution of this analysis.

Another DS of interest is Mokosh, which degrades foreign viral transcripts to protect cells from infections^8^. Mokosh system proteins (MkoA and MkoB) have domain homology to the anti-transposon piRNA (PIWI-interacting RNA) pathway in eukaryotes^5^. Our phylogenetic analysis of MkoA suggests a bacterial origin of this protein. However, MkoA homologs associated with defense functions in eukaryotes are distributed in other regions of the phylogenetic tree. Thus, further analyses are needed to elucidate the relationship between functional eukaryotic immune proteins and MkoA homologs (Supplementary Figure 2B). Like much of the history of cellular complexity in eukaryotes, these findings suggest that eukaryotic DS were derived from both bacteria and archaea (Figure 4).

**Figure 4.**
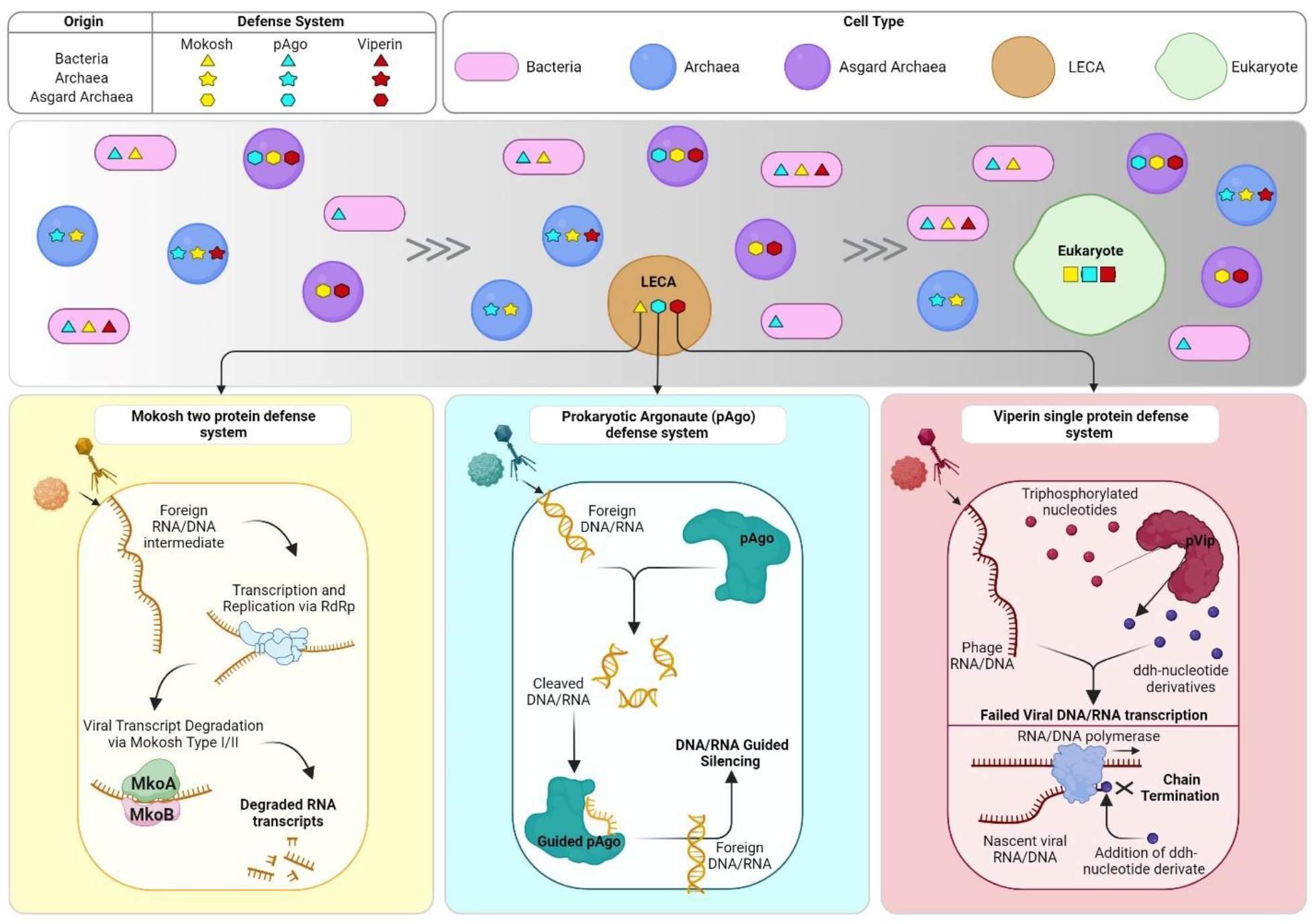
Model illustrating the contributions of archaea to the origins of eukaryotic defense mechanisms. Based on our reconstructions of structures and phylogeny in the study, we propose that early forms of both the viperin and argonautes defense systems were inherited from Asgardarchaeota via the LECA (last eukaryotic common ancestor) into modern eukaryotes. However, other systems present in Asgard like mokosh appear to have originated from bacteria, highlighting that both prokaryotes have an important role on the origin of defense mechanisms in Eukaryotes. The known mechanisms of mokosh, argonaute, and viperin defense systems are illustrated at the bottom. Created using BioRender.com.

This study sheds light on the roles of archaea in the origin of innate immune mechanisms in eukaryotes. Asgardarchaeota emerges as a key player in the origins of viperins and argonautes, with ancestral forms of these proteins tracing back to the last eukaryotic common ancestor (LECA) and prior (Figure 4). We found remarkable conservation observed in the sequences, structure, and function of these DS proteins compared to their eukaryotic counterparts. Models of eukaryogenesis involve the interaction between a bacterium and an archaeon; our findings reveal that early eukaryotes inherited robust defense mechanisms against viral infections from both ancestral partners (Figure 4). The compatibility of some of these proteins (asVips and asAgos) with those in eukaryotes suggests these systems may provide biomedical or biotechnology applications in eukaryotes.

## Methods

### Known Defense systems identification

A database of Archaeal genomes available from NCBI was built containing 646 genomes from the DPANN group, 668 genomes from TACK superphylum, 1408 genomes belonging to Euryarchaeota phylum, and 869 Asgard archaea genomes. This database only contains complete or high-quality genomes. To guarantee the quality of the genomes, a CheckM v1.2.1 screening was performed and only genomes presenting a completeness higher than 50%, and contamination lower than 10% were included. Another database containing 716 bacterial genomes from 71 different phyla was built following the same procedure. The distribution and diversity of known defense systems in archaeal genomes was checked running DefenseFinder v1.0.9^7^ with models v1.2.2. and default parameters on the custom databases.

### Selecting homologues of archaeal defense system proteins in Eukaryotic genomes

A database of defense system proteins homologues found in Eukaryotes created by Cury *et al*., 2022^5^ was used as a template to search for homologues of archaeal defense system associated proteins. A DIAMOND v2.0.13.151^36^ database was created with Cury’s dataset and 4 separated queries were prepared, one with proteins from each archaeal group (DPANN, Eury, TACK and Asgard) (Supplementary Data 1). A pairwise alignment was performed with DIAMOND v2.0.13.151 using BLASTP with a cutoff of >30% identity and <E^-05^ e-value. In addition, DefenseFinder was run on genomes from the EukProtV3 database^37^ to search for additional homologues of prokaryotic defense systems in eukaryotes following the same parameters described in the section above. In Figure 1A bubble plot, the less frequent DS in Eukaryotic genomes (31 in total: RTnitrilase-Tm, Old Tim, Hydrolase-3Tm, HelicaseDUF2290, gop beta cll, DUF4238, DprA-PPRT, 3HP, 2TM 1TM TIR, RosmerTA, PD-T7-5, PD-T7-1, PD-T4-3, PD-T4-2, PD-T4-1, Gao Tmm, Gao TerY, Gao RL, Gao Qat, Gao Ppl, Gao Mza, Gao let, Gao Hhe, Gao Her, RnlAB, Old exonuclease, Nhi, PD-T4-6, PD-T4-5, AbiT, AbiQ) were excluded from the visual representation, but can have their distribution check in each dataset on Supplementary Data 1.

### Phylogenetic analyses

The viperins phylogenetic analysis is shown in Figure 3. It contains viperin protein sequences identified by DefenseFinder in the archaeal database together with the viperins previously characterized and used on the phylogenetic analyses on Bernheim et al., 2021^13^ work. After removing redundant sequences with seqkit v.2.3.036, a total of 337 unique sequences were processed as described in the end of this section to generate a dendrogram (model selected: LG+R10).

To analyze the homology relationship between argonaute sequences, we combined the Long pAgo and eAgo sequences used by Swarts *et al*., 2014^33^ with COG1431_pAgo and pAgo_LongA sequences identified in our archaea database and the EukProtV3 database^37^ with DefenseFinder. Identical sequences were removed using seqkit v.2.3.0^38^, resulting in a total of 543 unique argonaute proteins. The sequences were aligned using MAFFT v7.457^39^. Regions in the alignment containing >90% gaps were removed using ClipKIT 1.3.0^40^. The tree was computed with IQTree v2.0.3 using 1000 bootstrapping replicates, and the best model generated (LG+F+R10) was selected according to the Bayesian information criterion (BIC)^41^. The same alignment, gap removal, and dendrogram generation procedures were used for all the protein sequence phylogenetic analyses in this section. All information about the protein sequences used in these analyses can be found in Supplementary Data 4. Alignments used to generate the phylogenetic analyses can be found at https://figshare.com/s/a027e2310360cf6f81ac.

### Structural homology analyses

To analyze the homology between proteins used as defense mechanisms in prokaryotes and eukaryotes, protein sequences associated with defense systems were submitted to structural modeling in ColabFold v1.5.2^42^. Multi-sequence alignments were performed using the mmseq2 mode, and AlphaFold2_ptm model. Three recycling steps were used to optimize the computational power and running time.

The predicted protein models were combined with reference structures of the respective proteins acquired from RCSB Protein Data Bank (RCSB). A multi-structural alignment (MSTA) of these structures was performed using the default parameters of mTM-align tool^43^ to build a dendrogram using IQTree v2.0.3 (models: LG+I+G4 for viperins; VT+R4 for argonautes).

The pairwise RMSD matrix obtained from mTM-aligns (Supplementary Data 3) was used to select the proteins suitable for the 3D reconstruction of their alignment using the Needleman-Wunsch algorithm and BLOSUM-62 matrix on ChimeraX software^44^.

When necessary, protein annotation was done using Interproscan v.5.31-70.0^45^ with the default parameters (Supplementary Data 4). Alignments used to generate the structural analyses can be found at https://figshare.com/s/a027e2310360cf6f81ac.

### Bacterial growth and phage propagation

*E. coli* strains (MG1655, Keio ΔiscR27, DH5α) were grown in LB or LB agar at 37^°^C. Whenever applicable, media were supplemented with chloramphenicol (25 µg mL^−1^) and/or kanamycin (50 µg mL^−1^) to maintain the plasmids. T7 phage was propagated on *E. coli* BL21s using the plate lysate method. Lysate titre was determined using the small drop plaque assay method as previously described^46^.

### Plasmid construction

All primers were purchased from IDT (Supplementary Data 2). asVip genes were synthesized by Twist Biosciences, and codon optimized where indicated. Genes were sub-cloned into an inducible expression vector using Golden Gate assembly. All plasmids were propagated in *E. coli* DH5α and then purified and transformed into the Keio ΔiscR27 strain to upregulate iron-sulfur cluster production, as described previously^13,47^.

### Plaque assays

Plaque assays were performed according to standard protocols^46^. Overnight cultures were diluted 1:100 in LB supplemented with 1.25 mM MgCl_2_, 1.25 mM CaCl_2_ and Isopropyl β-D-1-thiogalactopyranoside (IPTG; final concentration of 0.5 mM) for induction of asVip expression and grown to an OD_600_ ∼ 0.3. Bacteria from these outgrowth cultures were combined with serial dilutions of phage lysate and incubated at 37°C for 15 minutes before being mixed with 0.5% agar and plated on agar plates. After solidifying, the plates were incubated overnight at 37°C. Plaques were imaged using an Azure Biosystems 600 imaging system, and their areas were measured using FIJI’s ‘Analyze Particles’ plugin^48^. Three biological replicates were pooled for analysis. A two-tailed t-test was used to calculate statistical significance.

## Supporting information

Suppl_data_1-DefenseFinder

Suppl_data_2-asVip testing info

Suppl_data_3-Structural analyses info

## Contributions

P.L., B.J.B. and V.D.A designed the study.

B.J.B. and I.J.F. supervised and provided infrastructure to execute this study.

K.A. and V.D.A. performed metagenomic data assemblage and binning, providing MAGs for the study.

P.L., E.A.P., K.C., and V.D.A built and curated the datasets used in this study.

M.E.L., D.S., and I.J.F. were responsible for the design and execution of viperins experimental analyses.

P.L., E.A.P. and K.C performed phylogenetic analyses.

P.L. executed the protein structural homology analyses. P.L., M.E.L, and B.J.B wrote the manuscript.

All authors contributed to the final version of the manuscript and approved it before submission.

