## Supplementary figures for "Asgard archaea defense systems and their roles in the origin of eukaryotic immunity"

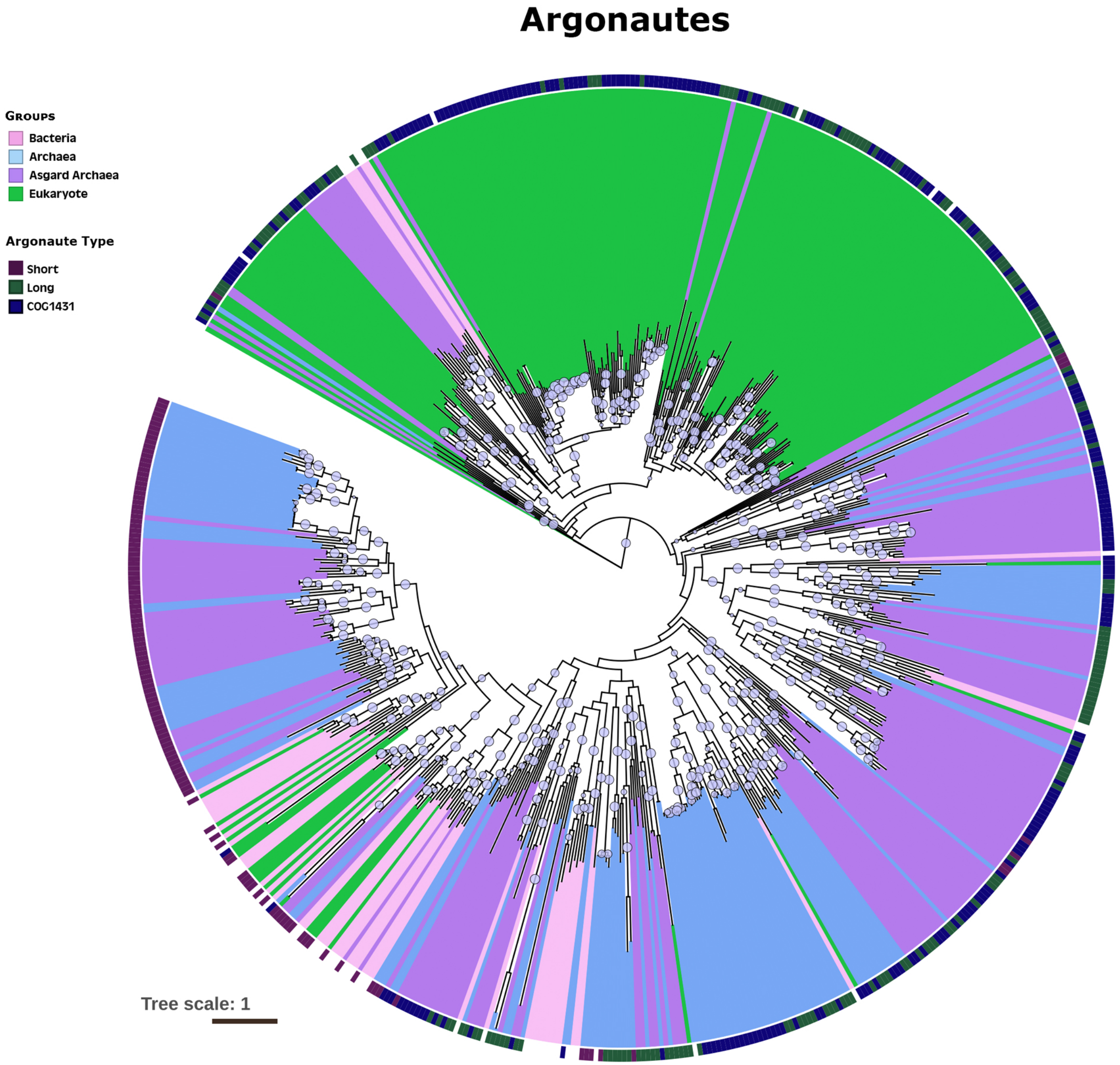


**Supplementary Figure 1**: Phylogenetic analyses of Argonaute proteins across the tree of life. Argonaute identify using DefenseFinder in each of the datasets prepared in this paper (See methods). A clear separation between argonautes from prokaryotic origin (bacterial (pink), archaeal (blue), asgard (purple)), and eukaryotic origin (green) can be observed. A clear segregation of the majority of short argonaute sequences (dark purple ring) is initially observed. Dots at the center of the branches represent Boostsrap values between 70-100.

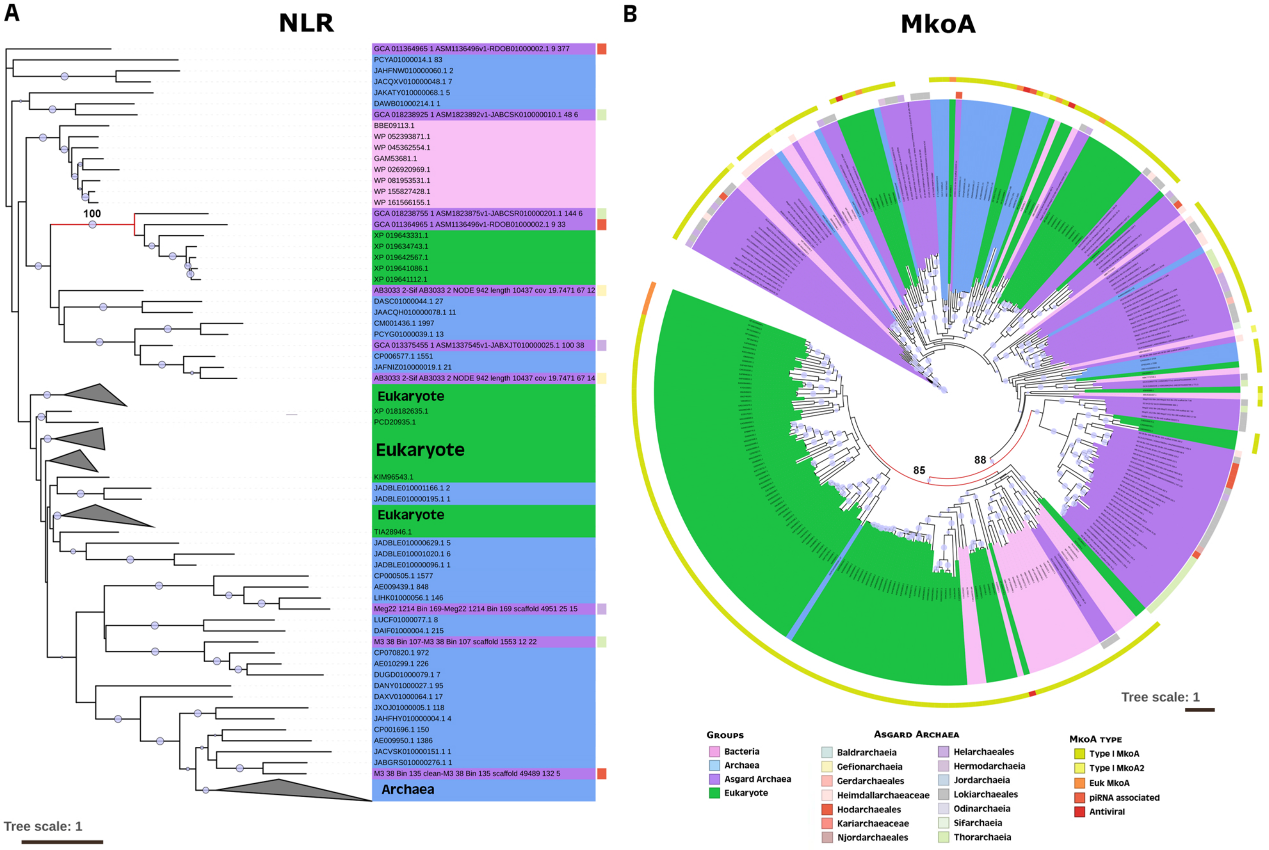


**Supplementary Figure 2**: Phylogenetic analyses of prokaryotic Defense system proteins and their homologues in Eukaryotes. A) Prokaryotic NLR proteins, and eukaryotic homologues. A preliminary correlation between Archaeal NLR and their eukaryotic counterparts is observed. Especially between a small group of Eukaryotes and asgard NLR sequences (red node). B) The Mokosh defense system proteins MkoA shows a vast array of homologues in eukaryotic genomes that appear to be from prokaryotic origin (red nodes). The eukaryotic homologues linked before to immune mechanisms in a previous study[5] (red on outer ring) are closer to more ancient versions of prokaryotic proteins and widespread in the tree. Dots at the center of the branches represent Boostsrap values between 70-100
